# Astrovirus in Reunion Free-tailed Bat (*Mormopterus francoismoutoui*)

**DOI:** 10.1101/774224

**Authors:** Léa Joffrin, Axel O. G. Hoarau, Erwan Lagadec, Marie Köster, Riana V. Ramanantsalama, Patrick Mavingui, Camille Lebarbenchon

## Abstract

Astroviruses (AstVs) are RNA viruses infecting of a large diversity of avian and mammalian species, including bats, livestock, and humans. We investigated AstV infection in a free-tailed bat species, *Mormopterus francoismoutoui*, endemic to Reunion Island. A total of 380 guano samples were collected in a maternity colony during 38 different sampling sessions, from June 21^st^ 2016 to September 4^th^ 2018. Each sample was tested for the presence of the AstV RNA-dependent RNA-polymerase (RdRp) gene using a pan-AstV semi-nested polymerase chain reaction assay. In total, 27 guano samples (7.1%) tested positive, with high genetic diversity of the partial RdRp gene sequences among positive samples. A phylogenetic analysis further revealed that the detected viruses were genetically related to AstVs reported in rats, reptiles, dogs, and pigs, but did not cluster with AstVs commonly found in bats. Although more investigations need to be conducted to assess the prevalence of infected bats in the studied population, our findings show that Reunion free-tailed bats are exposed to AstVs, and suggest that cross-species transmission may occur with other hosts sharing the same habitat.

## Introduction

Astroviruses (AstVs) are positive-sense single-stranded RNA viruses and have been detected in a large diversity of mammalian and avian species (De Benedictis, Schultz-Cherry, Burnham, & Cattoli, 2011). In humans, eight serotypes have been described worldwide, accounting for 2%–9% of acute non-bacterial gastro-enteritis cases in children (Bosch, Pinto, & Guix, 2014). AstVs are also responsible for diseases in livestock, poultry and domestic pets (De Benedictis et al., 2011). In wild animals, they have been mostly described in bats (Fischer, dos Reis, & Balkema-Buschmann, 2017) and in aquatic birds (Chu et al., 2012), although detection in other hosts has been reported, such as marine mammals (Rivera, Nollens, Venn-Watson, Gulland, & Wellehan, 2010) and non-human primates (Karlsson et al., 2015). In addition to this large host diversity, the evolutionary history of AstVs is characterized by frequent cross-species transmission events (Mendenhall, Smith, & Dhanasekaran, 2015), supporting limited host specificity and a risk of spillover and emergence in humans (Bosch et al., 2014).

AstVs have been reported in African insectivorous bats, in particular in species of the Miniopteridae and Rhinonycteridae families (Hoarau et al., 2018; Rougeron et al., 2016; Waruhiu et al., 2017; Yinda et al., 2018). Detection rates vary greatly depending on the study design and the tested species (Fischer et al., 2017), and the prevalence of bat shedding AstVs within populations is also highly variable. For instance, we estimated prevalence of shedding bats ranging from 20% to 81% for *Triaenops afer* in Mozambique, depending on the time of sample collection (Hoarau et al., 2018). Such a strong seasonality has been reported in both tropical and temperate regions and could be associated to bat population dynamics, in particular in maternity colonies (*e.g*. increase in the population size and density during the formation of the colony; Drexler et al., 2011). Body condition and co-infection with other viruses have also been identified to be positively correlated with AstV infection in bats (Mendenhall et al., 2017; Seltmann et al., 2017).

The islands of the Western Indian Ocean host eleven of the twenty bat families described worldwide (Goodman, 2011), with a highly contrasted diversity pattern between islands. Madagascar shelters 46 bat species, of which nearly 80% are endemic (Goodman et al., 2016; Foley et al., 2017). In contrast, the small oceanic islands surrounding Madagascar are inhabited by a limited number of bat species. For instance, only three species occur on Reunion Island and only one of them, the Reunion free-tailed bat (RFTB; *Mormopterus francoismoutoui*; Molossidae), is endemic to the island.

RFTB is the most abundant bat species on Reunion Island. This small insectivorous species roosts in large colonies in natural caves and synanthropic habitats such as bridges and houses (Goodman et al., 2008). Based on guano sampling in a RFTB maternity colony, we investigated the temporal variation in AstV shedding for more than 2 years (28 consecutive months), with a particular emphasis on the parturition period. Partial sequencing of the AstV RNA-dependent RNA-polymerase (RdRp) gene and phylogenetic analyses were performed to assess the diversity and the origin of the detected viruses.

## Material and methods

The study was conducted in the main RFTB colony known to date on Reunion island, within a natural cave of 30 m^3^ in the West coast of the island (21,111°S, 55,259°E). The colony is monospecific and mostly composed of adult females during the early stages of the breeding season (October to December); and of neonates and juveniles, after the parturition period (from mid-December onward; Dietrich et al., 2015). Between 40 000 and 50 000 flying adult bats have been estimated before parturition; however, after the breeding season, the cave remains totally empty from bats (May to September) (Dietrich et al., 2015).

A non-invasive sampling was setup to limit colony disturbance. During each sampling session, we collected guano samples in the narrowest part of the cave entrance, in ten different sampling points separated by 50 cm along a 4.5 m transect (Supplementary Figures 1 and 2). Sterile open-ended 2 mL syringes were used to collect a core sample, in order to obtain *c*. 130 mg of guano. Each guano sample was immediately mixed with 1.5 mL of Virus Transport Media (VTM; *e.g*. Lebarbenchon et al., 2017). Samples were maintained in a cooler with ice packs in the field and stored in a −80°C freezer within two hours.

In total, 380 samples were collected during 38 different sampling sessions between June 21^st^ 2016 and September 4^th^ 2018. During each sampling session, the colony was visually monitored in order to classify the relative population size (no bat, less than half of the cave covered with bats, more than half of the cave covered), and to assess the population age structure and timing of parturition based on the presence of adults, neonates, and juveniles.

Guano samples were mixed and centrifuged at 1 500 g for 15 min. RNA extraction was performed on 140 μL of supernatant, with the QIAamp Viral RNA Mini Kit (QIAGEN, Valencia, CA, USA).

Reverse-transcription was done on 10 μL of RNA using the ProtoScript II Reverse Transcriptase and Random Primer 6 (New England BioLabs, Ipswich, MA, USA) (Lebarbenchon et al., 2017). cDNAs were tested for the presence of the AstV RdRp gene using a pan-AstV semi-nested polymerase chain reaction (PCR) assay (Chu, Poon, Guan, & Peiris, 2008). PCR products of the expected size were submitted for direct Sanger sequencing (Genoscreen, Lille, France). Chi-square tests were conducted to investigate the effects of the sampling date and of the sampling location in the transect, on the probability of successful detection of AstV RdRp gene. Statistical analyses were conducted with R 3.4.4 (R Core Team, 2018).

A phylogenetic analysis was performed on the RdRp gene sequences obtained in this study (GenBank accession numbers MK966904 to MK966918, and MZ274075 to MZ274086) and 84 reference RdRp sequences of AstVs detected in a large diversity of bat species, in particular from Africa (Madagascar, Mozambique, Gabon), as well as in other vertebrate hosts (*e.g*. avian, human, livestock). Sequences were aligned with CLC Sequence Viewer 8.0 (CLC Bio, Aarhus, Denmark). A maximum-likelihood analysis was done using PhyML 3.1 (Guindon et al., 2010), with an evolutionary model selected by Model Generator 0.85 (GTR + I + Γ; I = 0.13, α = 0.83; Keane, Creevey, Pentony, Naughton, & Mclnerney, 2006), and 1 000 bootstraps.

## Results and discussion

In total, 27 guano samples (7.1 %) tested positive for the presence of AstV, without difference between sampling location along the transect (*χ*^2^ = 11.8; df = 9; *P* = 0.23). Although no variation was observed between the two sampling seasons (*χ*^2^ = 0.17; df = 1; *P* = 0.68), significant differences in the number of positive samples between sampling dates were detected (*χ*^2^ = 74.8; df = 37; *P* < 0.001), with AstVs mostly detected after the parturition period (late December), in February and March (Figure 1). This pattern suggests a potential effect of the population age structure (Mendenhall et al., 2017) rather than changes in size and density associated to the formation of the colony (Drexler et al., 2011). A previous investigation on paramyxovirus and *Leptospira spp*. infection dynamics in the same bat colony revealed two peaks of infection: one during the colony formation and the other two months after the birth pulse (Dietrich et al., 2015). The later may coincide with the detection of most AstVs in February and March, and the first one with the positive samples detected in December. Although these results support the occurrence of AstV RNA in the RFTB environment, our sampling design precludes further conclusion on the prevalence of shedding bats in the population. Bat individual sampling is needed to further support AstV infection in RFTB and identify population-related factors involved in the temporal and spatial transmission dynamics of AstVs.

**Figure 1.**
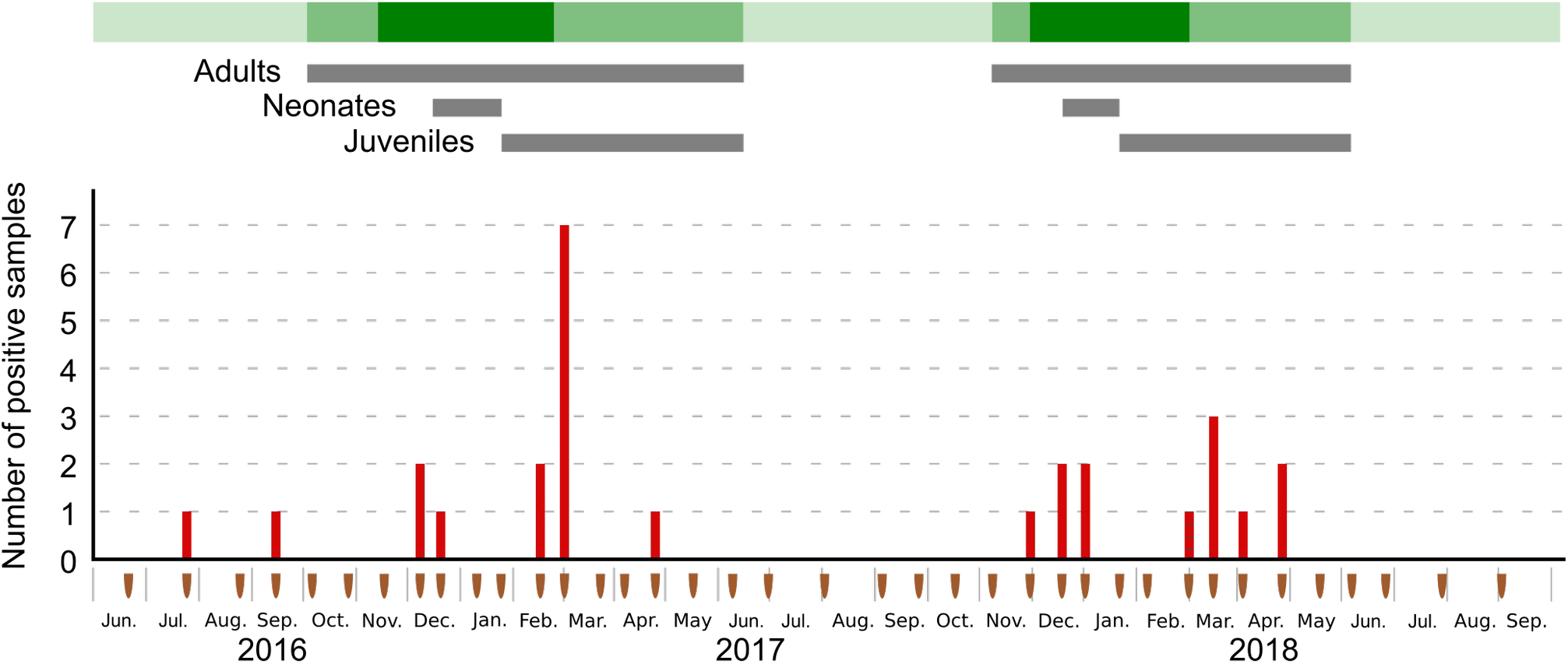
Number of Astrovirus positive guano samples as per sampling session (brown marks). Green shading indicates the relative bat population size: light green: no bat inside the cave, mid-green: less than half of the cave occupied by bats, dark green: more than half of the cave occupied by bats. Presence of adults, neonates, and juveniles bats is represented by gray bars.

Two samples collected before the breeding season (in July and September 2016), when the cave was not occupied by bats, also tested positive for AstV, and suggest a possible environmental persistence of AstV RNA in bat guano. AstV persistence in the environment could potentially favor bat infection during cave colonization at the beginning of the breeding season, but also create opportunity for virus spillover to other vertebrate hosts. Rats (*Rattus rattus*) have been sighted at the vicinity and within the cave during the study period, and previous studies have shown that paramyxoviruses may be transmitted between rats and RFTB in the studied cave (Dietrich et al., 2015). Bat guano is also used as a fertilizer for agricultural crops, worldwide, and is regularly harvested on a small scale by local farmers for personal use (Mildenstein & de Jong, 2011). At our study site, bat guano was indeed harvested in the cave entrance throughout the course of the breeding season. The detection of viral RNA does not provide evidence of the maintenance of infectious particles in bat guano and virus isolation would be required to experimentally assess the infectious ability of bat AstVs outside their hosts. However, transmission routes of bat viruses to humans remain unclear (Joffrin, Dietrich, Mavingui, & Lebarbenchon, 2018) and given the high propensity of AstVs for cross-species transmission (Mendenhall et al., 2015), our finding underlines that the risk of AstV spillover from bats to human should be fully investigated.

Analysis of the 27 RdRp partial sequences revealed a high genetic diversity (up to 49% of partial RdRp sequence differences). This finding is consistent with previous studies on bat AstVs in Madagascar (Lebarbenchon et al., 2017), continental Africa (Hoarau et al., 2018; Rougeron et al., 2016; Waruhiu et al., 2017; Yinda et al., 2018), and other bat populations, worldwide. Such a high level of sequence diversity in a single bat species nevertheless contrasts with the low diversity measured for paramyxovirus and *Leptospira* in the same colony (Dietrich et al., 2015). Classification of AstV based on host species origin has been largely challenged for the past years (Donato & Vijaykrishna, 2017), because AstVs isolated from different animal species can be genetically similar (Hoarau et al., 2018; Lebarbenchon et al., 2017; Waruhiu et al., 2017), but also because genetically different viruses can be isolated from a single species (Waruhiu et al., 2017; Yi et al., 2018). To improve AstVs classification, the International Committee on Taxonomy of Virus now recommends to compare AstV RdRp gene to capsid gene sequences (Bosch et al., 2011).

The phylogenetic analysis further revealed that the detected viruses were genetically related to AstVs found in rats, reptiles, dogs, cats and pigs, but did not cluster in a single bat AstVs clade (Figure 2). This result could reflect the limited number of AstV nucleotide sequences available from Molossid bats in public database and, more broadly, from bats in tropical areas. Because our study was based on environmental sampling, however, we cannot exclude that the detected viruses came from other hosts than RFTB. As previously mentioned, rats, but also feral dogs (*Canis lupus familiaris*), and zebu (*Bos taurus indicus*), have been regularly sighted close to the cave entrance. Avian species such as house sparrow (*Passer domesticus*), Madagascar turtle-dove (*Nesoenas picturatus*), and common myna (*Acridotheres tristis*) have also been observed near to the cave, and can be in direct contact with bat guano. Further investigations involving sampling of individual bats and other host species in the vicinity of the cave, as well as sequencing of the AstV capsid gene, could provide information on the level of potential cross-species transmission in this ecosystem.

**Figure 2.**
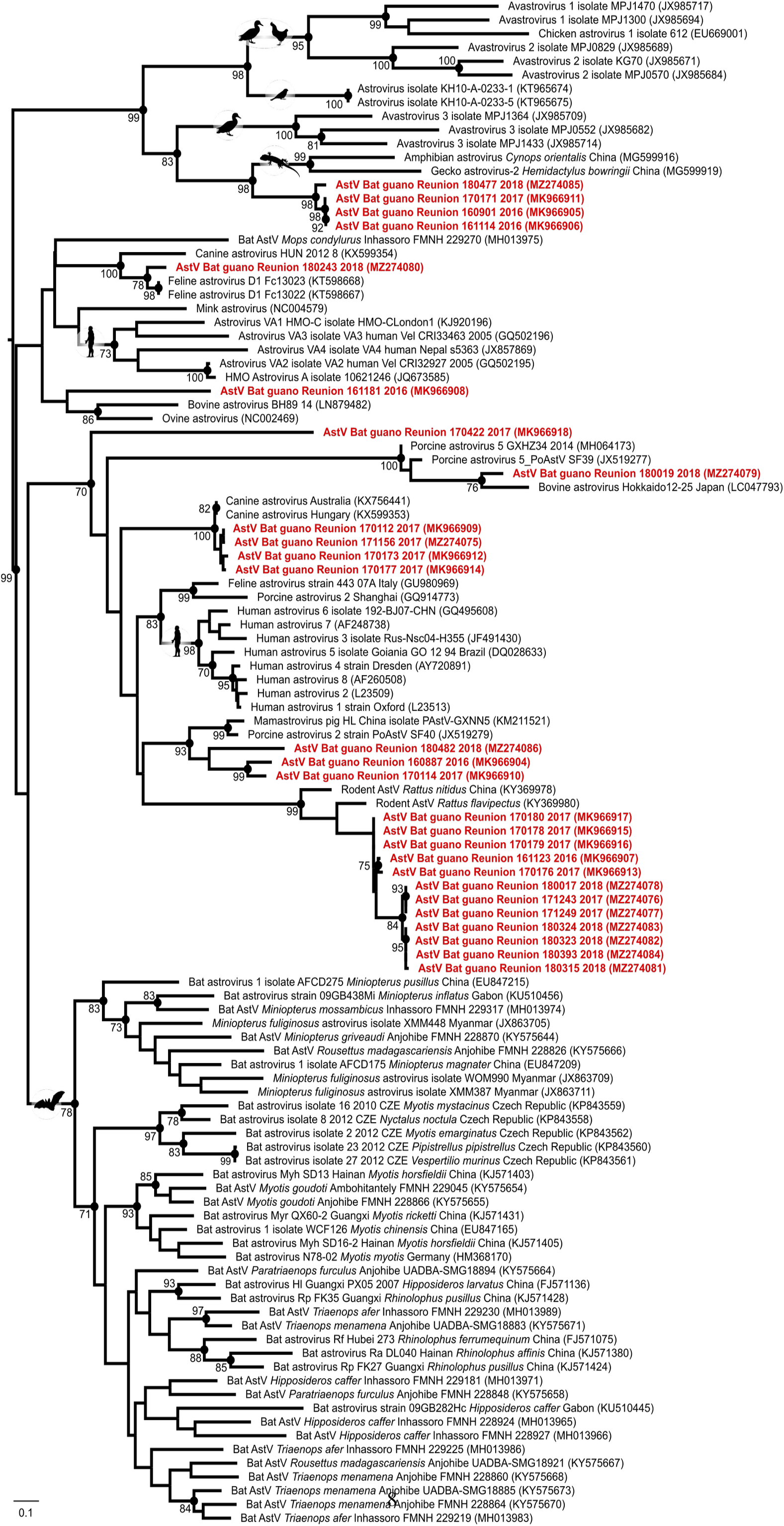
Maximum likelihood consensus tree derived from 111 Astrovirus (AstV) RNA-dependent RNA-polymerase partial nucleotide sequences (390 bp). Black dots indicate nodes with bootstrap values higher or equal than 70. Red: sequences generated in this study. Scale bar: mean number of nucleotide substitutions per site.

## Conflicts of interests

The authors have no conflict of interests.

## Acknowledgments

We thank Muriel Dietrich and Nicolas Diotel for their assistance in the field. Léa Joffrin was supported by a “Région Réunion, European Regional Development Funds (FEDER 2014-2020)” PhD fellowship, Axel Hoarau by a “Ministère de l’Enseignement supérieur, de la Recherche et de l’Innovation” PhD fellowship, and Camille Lebarbenchon by a “Chaire Mixte Institut National de la Santé et de la Recherche Médicale (INSERM) - Université de La Réunion”. This study was funded by the VIROPTERE program (INTERREG V Océan Indien).

## Notes

### Competing Interest Statement

The authors have declared no competing interest.

