## Supplementary Figures for "Astrovirus in Reunion Free-tailed Bat (*Mormopterus francoismoutoui*)"

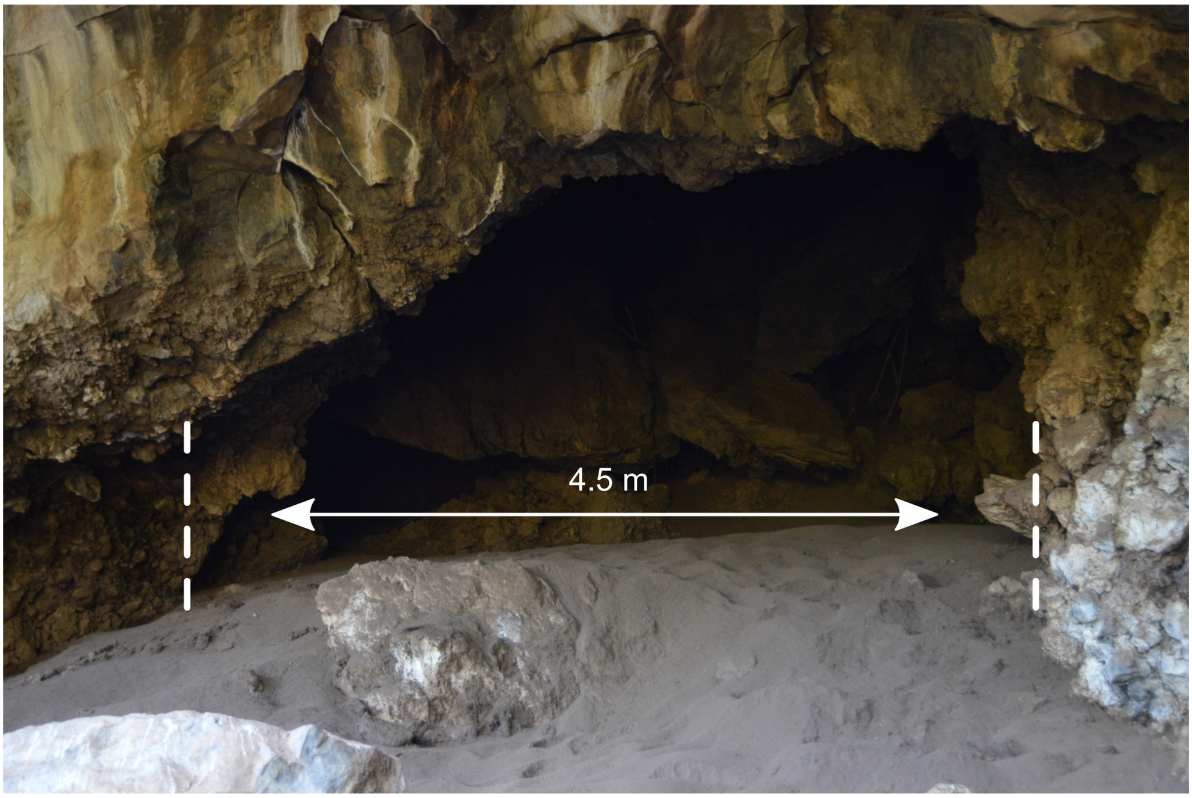


**Figure S1.** Cave entrance of the studied colony. The narrowest part of the entrance is approximately 4.5 m width and 2m high.

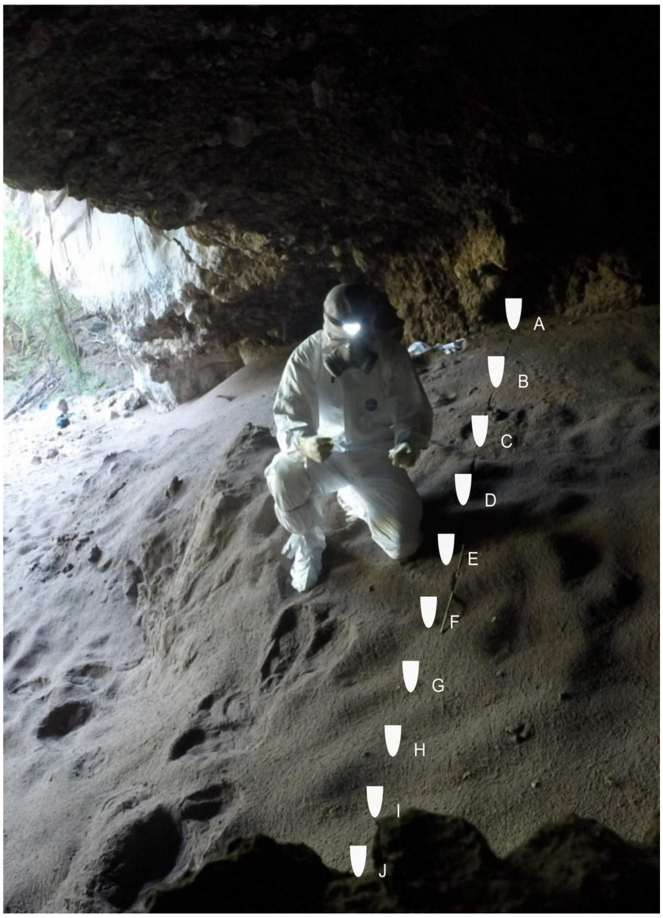


**Figure S2.** Sampling transect at the cave entrance. Ten guano samples separated by 50 cm (A to J) were collected at each sampling session.
